# Life-course social disparities in body mass index trajectories across adulthood: cohort study evidence from China Health and Nutrition Survey

**DOI:** 10.1101/2022.08.23.505012

**Authors:** Yusong Dang, Peixi Rong, Xinyu Duan, Mingxin Yan, Yaling Zhao, Baibing Mi, Jing Zhou, Yulong Chen, Duolao Wang, Leilei Pei

**Author notes:** Corresponding author: Leilei Pei, Department of Epidemiology and Health Statistics, School of Public Health, Xi’an Jiaotong University Health Science Center, Xi’an, Shaanxi 710061, P.R. China.

## Abstract

**Background:** The social disparities in obesity may originate in early life and adult social class. There are various developmental trajectories of overweight/obesity in adulthood. It is unclear how the intergenerational mobility of socioeconomic status influences adult overweight/obesity in China.

**Methods:** We used longitudinal data from ten waves of the China Health and Nutrition Survey (CHNS) between 1989 and 2015 for our analysis. The group-based trajectory modeling was used to identify BMI trajectories in adulthood. Multinomial logistic regression was adopted to assess the associations between SES and adult BMI trajectories.

**Results:** Among a total of 3,138 participants, three latent clusters, including normal-stable BMI (51.4%), progressive overweight group (39.8%), and progressive obesity group (8.8%), were identified. High father’s occupational position, high participants’ occupation position and educational attainment, respectively, were associated with greater obesity risk. Compared to a stable low life course SES trajectory, a stable high life course SES trajectory was associated with a 2.35-fold risk of obesity, and upward and downward social mobility trajectories increased the risk for overweight/obesity. Individuals in the highest relative to the lowest life course cumulative socioeconomic score group had around twice risk of obesity.

**Conclusions:** The results emphasize the role of the high SES in early life and life-course SES accumulation, in the obesity intervention in China.

**Funding:** All the work was supported by the National Natural Science Foundation of China (Grant Nos. 72174167, 81602928) and Natural Science Foundation of Shaanxi (2021JM-031).

## Introduction

Obesity has recently been a public health concern around the world, due to its strong association with many negative physical and mental health outcomes, including hypertension, diabetes, cancer and depression as well as mortality(1–3). Recent studies suggest that in China between 2015 and 2019, the prevalence of overweight and obesity in adults have reached 34.3% and 16.4%, respectively(4). The obesity and overweight epidemic have been influenced by both personal and environmental factors, such as genes, diet culture and behavioral patterns(5). The bulk of evidences show that socioeconomic status (SES), such as education, occupation and income, very often lead to social differences in the prevalence of obesity and overweight(6, 7). The majority of studies from western countries find an inverse relationship between SES and obesity(8, 9). In the developing countries, this interrelationship shows a strong positive correlation between those. Previous study in Chinese juvenile and adult population found higher income contributed to the risk for obesity(10, 11). With the rapid development of China’s economy, the relationship between SES and obesity is becoming increasingly complex. Hence, understanding SES disparities in obesity is presently an essential element for Chinese government in establishing public health priorities.

Socioeconomic disparities seem to have their origins in early life and perhaps even in earlier generations. In the United States, for example, a longitudinal 19-year study suggests that higher SES in childhood protects against weight gain at age 18 and in adulthood(12). Furthermore, it is now well established that adults suffer from gain weight and excess adiposity mostly in this period from young to middle adulthood, while weight remain fairly stable from middle to late adulthood, suggesting that there are various developmental trajectories of overweight/obesity in adulthood(13). In developing countries, however, studies on the relationship between the SES in early life and adult BMI trajectory are scarce, and it is also unclear whether early life course SES still has an impact on adult BMI. Therefore, it is necessary to investigate the developmental characteristics of adult overweight/obesity across different SES in developing countries, particularly across early life SES trajectory, to provide measures to reduce future disease burdens.

In this study, we extracted data from a 20-year longitudinal survey, the China Health and Nutrition Survey (CHNS), to assess the influence of life-course SES on adult obesity development. Our main objectives are to evaluate: (1) latent effects of early life socioeconomic circumstances on adult obesity; (2) the effect of SES over the life course on BMI trajectory in adulthood.

## Methods

### Study population

The data was extracted from China Health and Nutrition Survey (CHNS), which is an ongoing cohort study to explore how a series of economic, sociological, and demographic factors influence health and nutritional status of interest in China’s population. The first wave of CHNS began in 1989, and subsequently the surveys were tracked in 1989, 1991, 1993, 1997, 2000, 2004, 2006, 2009, 2011 and 2015. A multistage random cluster sampling method was adopted in the survey, covering 239 communities from nine of China’s 31 provinces. The survey was approved by institutional review boards at the University of North Carolina, Chapel Hill (Chapel Hill, NC), and China Center for Disease Control and Prevention (Beijing, China), and each participant provided written informed consent. A more detailed description of the design and procedures of CHNS has been described elsewhere(14, 15).

We used longitudinal data from ten waves of the CHNS between 1989 and 2015 for our analysis. The flow diagram of the study cohort is summarized in Figure 1. Initially, a total of 38,536 participants were extracted from the original surveys. Then approximately 35398 participants were excluded due to the following reasons: < 18 years old (n=7,870), less than 2 visits of BMI in adulthood (n=11,768), missing father’s occupational position (n=15,604), missing education (n=29) and economic status in adulthood (n=92), and pregnant women (n=35). Finally, a total of 3,138 participants with 11440 visits were included in the study.

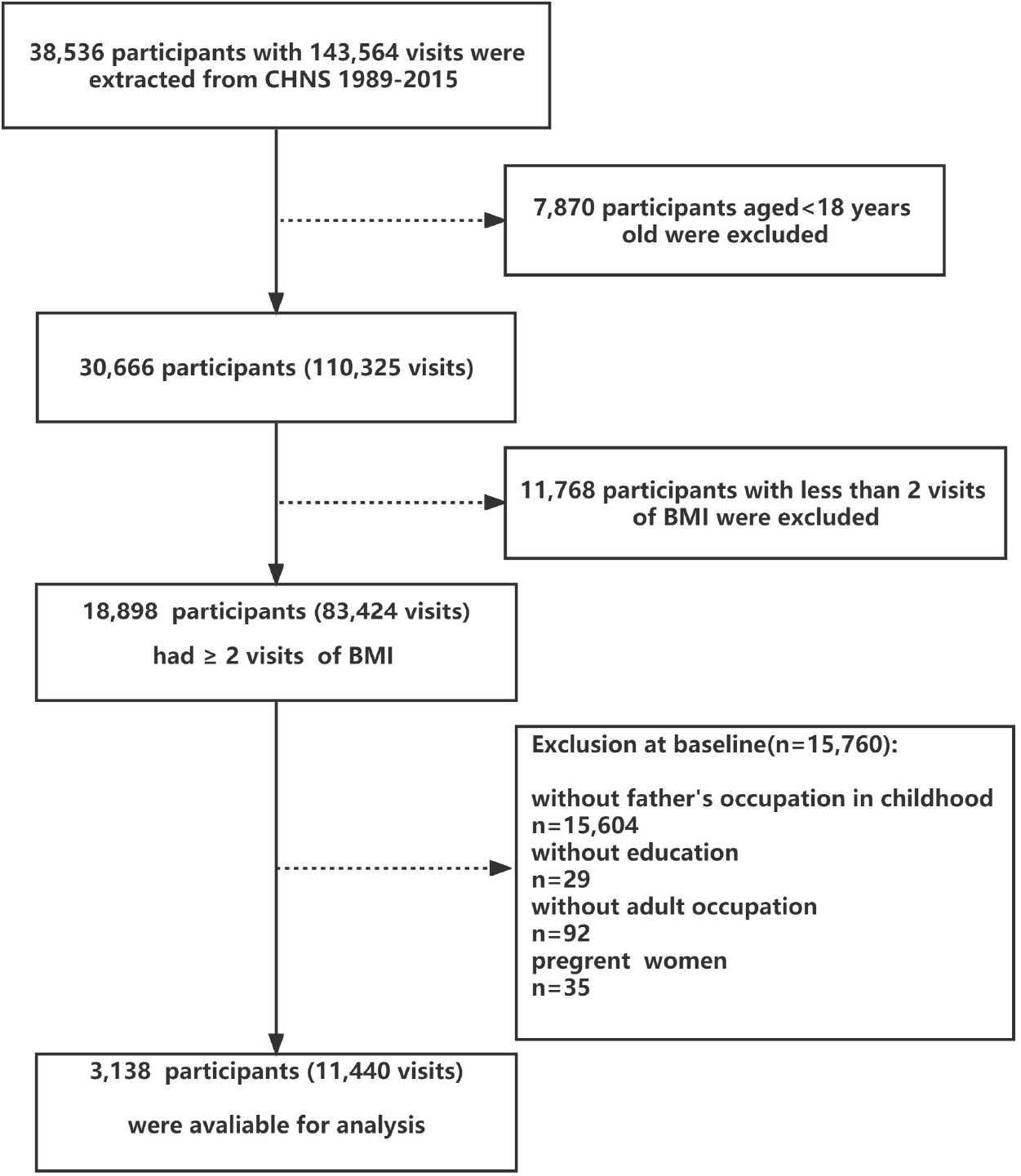

### Outcome assessment

In the study, the weight was measured using a balance-beam scale with an accurate to 0.1 kg, when participants wore only light clothing. The height measurement was conducted with participants barefoot on a portable stadiometer that was accurate to 0.1 cm. BMI was calculated as weight (in kilograms) divided by the square of height (in meters). According to the Working Group on Obesity in China (WGOC), the BMI based definition of overweight or obesity for the Chinese population is lower than that for the European or North American population. In the study, thus, overweight and obesity were defined as BMI ≥24 kg/m^2^ and BMI ≥28 kg/m^2^, respectively(16). To represent the development of BMI in adults, all BMI measurements from 1989 to 2015 were utilized to explore BMI trajectories with the age in the survey year as the timeline.

Group-based trajectory model (GBTM) was adopted to analyze the dynamic changes of adult BMI during follow-up and identify potential clusters with similar trajectories(17). These trajectories were summarized by a finite set of different polynomial functions of age or time, determining the form and number of groups that best fit the data. Briefly, GBTM provides a statistical method to identify the trajectory of each group and the form of each trajectory, and estimates the probability for each individual of group membership and assign them to one of the trajectories for which they have the highest estimated probability. GBTMs with different numbers of latent classes were compared using information criteria (IC)-based fit statistics, including Akaike Information Criteria (AIC), Bayesian Information Criteria (BIC), Log-Likelihood G2. Lower values on these fit statistics were used for the best-fitting model selection. The value of average posterior probability (entropy) was commonly used to assess whether individuals are accurately classified into their most likely class, of which the value ranges from 0 to 1, with scores higher than 0.7 reflecting classification more accurate. At the same time, each latent class with a reasonable sample size was also considered for model selection, which will make sure interpretability of results. GBTM was conducted using Stata 12.0 with the traj procedure.

### Socioeconomic indicators

We collected three indicators at the baseline survey, including father’s occupation position, the participant educational attainment and adult occupation position, to represent individual socioeconomic status over the life course. Father’s occupational position commonly indicated SES of participants in childhood, which was assessed retrospectively with the question “What is/was your father’s main job”. Similarly, the participant occupational position in adulthood was acquired by the question “What is/was your main job”, which is the most used indicators of adult SES(18). According to the classification criteria of socioeconomic classification scheme(19), father’s and participant’s occupational positions were categorized into high (social classes I–Π), medium (social classes III–IV) and low (social class V) (Table S1, Supplementary file 1), coded from 0 (low) to 2 (high).

The participant educational attainment was considered as a main indicator of SES in his own early life, especially in young adulthood(6). Education was defined as the highest qualification in full-time education, was obtained by the question “How many years of formal education did you have in formal school?” In the study, education was also grouped into three categories: high (≽ 12 years formal education), medium (8-11 years formal education), and low (<8 years formal education).

Two indicators were established to represent participants’ life-course SES, including life-course socioeconomic trajectories and a cumulative SES score. The cumulative life-course SES score was sum of paternal occupational position and participant’s education and adult occupational position, ranging from 0 to 6, with higher values corresponding to greater life-course advantage. The life-course socioeconomic trajectories from childhood to adulthood were computed using information on the father’s occupational position and adult occupational position, both of them were dichotomized as high (social class I-IV) and low (social class V). Four possible combinations of socioeconomic trajectories across the life course were generated (Table S2): high SES in childhood and high SES in adulthood (stable high, n = 1524), low SES in childhood and high SES in adulthood (upward, n =581), high SES in childhood and low SES in adulthood (downward, n = 292), and low SES in childhood and low SES in adulthood (stable low, n = 741).

### Covariates

According to previous studies and a priori knowledge about our data(20), a set of covariates were considered as potential confounders, including gender, age, place of residence, smoking/drinking habits, total daily energy intake (TDEI) and physical activity level (PAL) were used as covariates in this study. Place of residence was determined by the question “Where do you live now, urban or rural?”, and was further classified into two groups (urban or rural). Smoking was obtained by the question “Have you ever smoked cigarettes (including hand-rolled or device-rolled)?” and were divided into two categories, “yes” (smoking currently) or “no” (not smoking currently). Drinking was acquired by the question “How often do you drink?”, and was also categorized into “yes” (drinking ≽ 1 per month last year) or “no” (drinking < 1 per month).

The total dietary energy intake (TDEI) was assessed at the household and individual levels, with individual’s intake collected using a weighing method in combination with three consecutive 24-h recalls (on 2 weekdays and 1 weekend day). TDEI was calculated based on the China Food Composition Table(21), which was used to determine the energy content in each food item consumed by the individual. The caloric value of all food items was summed to calculate a daily total, finally taking the average of total energy intake for 3 days. Physical activity level was derived in the survey, including very light PAL (sitting at work, such as office workers, watch repairers, etc.), light PAL (standing at work, such as sales clerks, laboratory technicians, teachers, etc.), moderate PAL (students, drivers, electricians, metal fabrication workers, etc.), heavy PAL (farmers, dancers, steel workers, athletes, etc.) and very heavy PAL (loaders, lumberjacks, miners, masons, etc.). PAL was reclassified into 3 groups: very light and light PAL activities were combined into light PAL group, moderate PAL included moderate PAL activities, and heavy and very heavy PAL activities was combined into heavy PAL group.

### Statistical analysis

The baseline characteristics were presented and compared across different socioeconomic status among participants. Continuous variables if normally distributed, were expressed as mean ± standard deviation and if not normally distributed, median and interquartile range was provided. Categorical variables were expressed as frequency and percentage. All univariate comparison of these variables across different subgroups was conducted by ANOVA, Wilcoxon rank-sum test, and Chi-square test.

A multinomial logistic regression model was used to explore the association between life-course socioeconomic status and adult BMI multi-class trajectories adjusting for some possible confounders. Adjusted odds ratios (OR) and 95% CI (confidence interval) were calculated for the different BMI trajectories. Each life-course SES indicator was first entered into a basic model, including gender, age and place of residence (Model 1). Smoking and drinking were further added in Model 2. Model 3 was further adjusted for variables in Model 2, plus physical activity level. Model 4 was further adjusted for variables in Model 3, plus total daily energy intake. Model 5 included all covariates from model 1 to model 4. Multicollinearity was checked using variance inflation factor (VIF). In the study, there are no variables with VIF > 4, representing no multicollinearity of the variables.

After the risk factors of interest were entered into Model 1 (gender, age and place of residence), the percent attenuation in the β coefficient for SES was calculated to express the contribution of risk factors in explaining the SES-BMI trajectory association. The formula was expressed as “100 × (β_Model 1_ — β_Model 1 + risk factors_)/(β_Model 1_)”. The 95% CI of the percentage attenuation was calculated using a bootstrap method with 1000 re-samplings.

A series of sensitivity analysis were conducted to test the robustness of the results. Firstly, we excluded participants who developed diabetes, hypertension, myocardial infarction, stroke or cancer at baseline from the sample, to minimize the possible reverse causation caused by these diseases. Secondly, we used the multiple imputation by the chained equations models to deal with missing covariates. We generated 20 imputed data sets, as the maximum missing amount of these variables was lower than 5%. The multinomial logistic regression analyses were repeated using each of the augmented data sets, and parameter estimates were averaged in the 20 sets. The adjusted OR values before and after multiple imputation were shown in the forest plot. All statistical analyses were performed with Stata version 12.0 and R version 4.1.3, and statistical significance was set at a 2-sided p < 0.05.

## RESULTS

### Long-term BMI trajectories across adulthood

The group-based trajectory models (GBTM) with 1-5 latent classes were fitted to select a reasonable model. When latent classes increased, AIC, BIC and Log-Likelihood G^2^ all continued to go down, and the decrease in the three-latent-class model leveled off (Figure S1, Supplementary file 1). Entropy was higher than 0.7 except for the four-latent-group model. Considering the interpretability of results, a frequency of >5% was preferred for each latent class (Table S3). Finally, the three-latent-class model was considered optimal (Figure 2): (1) normal-stable BMI (N=1613, 51.4%) with a consistently normal BMI, (2) progressive overweight (N=1249, 39.8%) with continuously increasing BMI from normal to overweight, and (3) progressive obesity (N= 276, 8.8%) with elevated BMI from overweight to obesity.

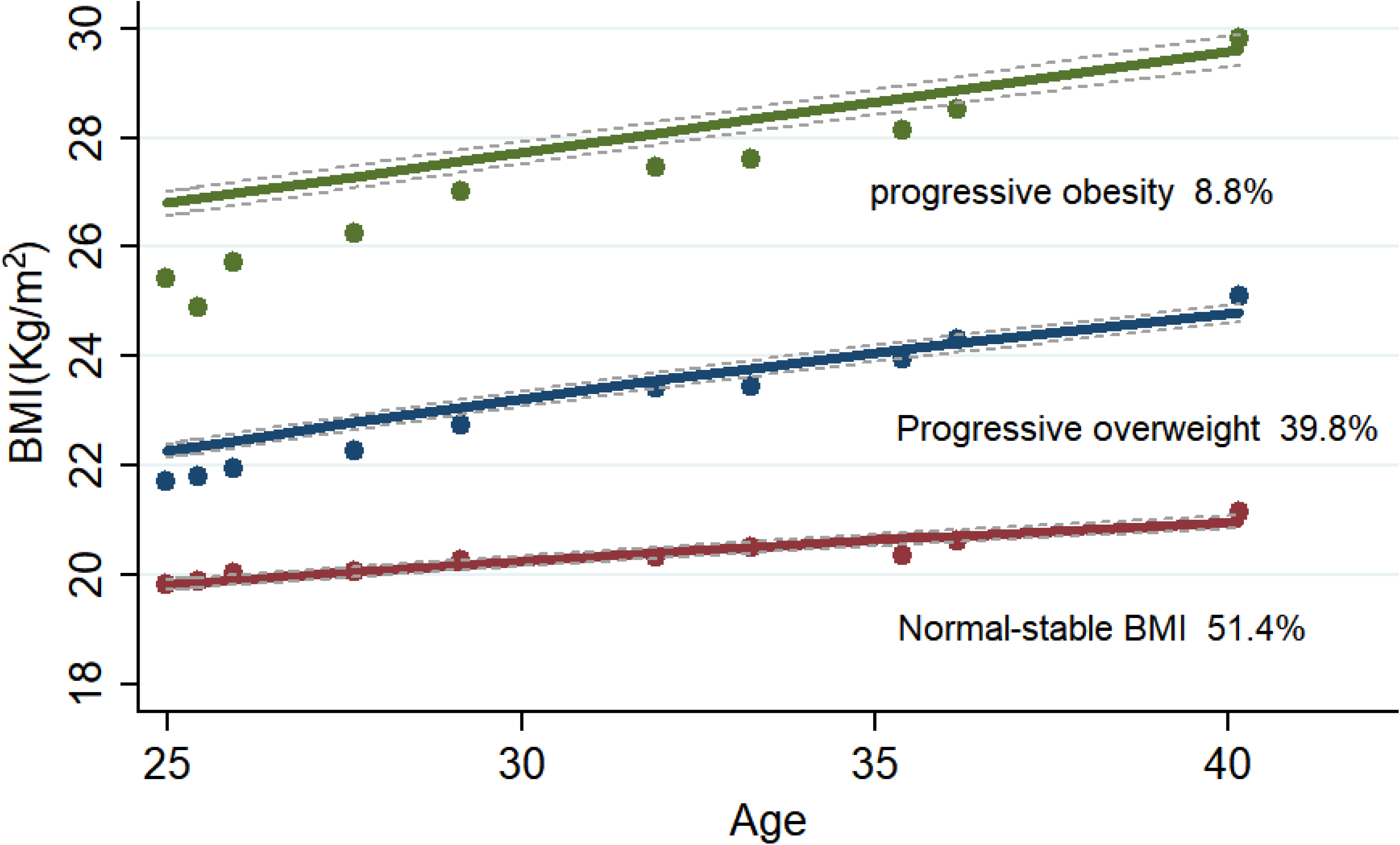

### Baseline characteristics of participants

Among 3,138 participants (909 women), the mean BMI of participants and education years were 21.2 ± 2.7 and 9.6 ± 3.2, respectively, and the median of age on years was 20(19–22) at baseline. The majority of participants (66.0%) lived in rural areas. Table 1 showed the baseline characteristics of the participants across different SES indicators.

**Table 1.** Baseline characteristics across socioeconomic status in early and adult life (n = 3138)^a^.

| Characteristics | Father's Occupational position |  |  |  | Adult's Education |  |  |  | Adult Occupational Position |  |  |  |
| --- | --- | --- | --- | --- | --- | --- | --- | --- | --- | --- | --- | --- |
|  | Low | Medium | High | *p* | Low | Medium | High | *p* | Low | Medium | High | *p* |
| Participants(n) | 1322(42.1) | 955(30.4) | 861(27.4) |  | 734(23.4) | 1398(44.6) | 1006(32.1) |  | 1033(32.9) | 1331(42.4) | 774(24.7) |  |
| Baseline Age(years) | 19.0(7.0) | 23.0(7.0) | 23.0(6.0) | 0.055 | 21.0(9.7) | 23.0(8.0) | 25.0(4.0) | <0.001 | 20.0(5.0) | 24.0(8.0) | 25.0(7.0) | <0.001 |
| Gender |  |  |  |  |  |  |  |  |  |  |  |  |
| Male | 1008(76.2) | 655(68.6) | 566(65.7) |  | 508(69.2) | 1024(73.2) | 697(69.3) | 0.049 | 711(68.8) | 996(74.8) | 522(67.4) | <0.001 |
| Female | 314(23.8) | 300(31.4) | 295(34.3) | <0.001 | 226(30.8) | 374(26.8) | 309(30.7) |  | 322(31.2) | 335(25.2) | 252(32.6) |  |
| Residential areas |  |  |  |  |  |  |  |  |  |  |  |  |
| Urban | 275(20.8) | 374(39.2) | 419(48.7) |  | 145(19.8) | 428(30.6) | 495(49.2) | <0.001 | 190(18.4) | 475(35.7) | 403(52.1) | <0.001 |
| Rural | 1047(79.2) | 581(60.8) | 442(55.1) | <0.001 | 589(80.2) | 970(69.4) | 511(50.8) |  | 843(81.6%) | 856(64.3) | 371(47.9) |  |
| BMI(Kg/m<sup>2</sup>) | 20.9±2.3 | 21.3±3.0 | 21.3±2.8 | 0.067 | 21.0±2.3 | 21.1±2.6 | 21.5±3.1 | 0.114 | 21.0±2.4 | 21.1±2.7 | 21.5±3.2 | 0.04 |
| *Health-related behaviors* |  |  |  |  |  |  |  |  |  |  |  |  |
| Smoking | 501(37.9) | 282(29.5) | 232(26.9) | <0.001 | 134(32.5) | 801(34.8) | 80(19.0) | <0.001 | 361(35.0) | 440(33.1) | 214(27.6) | 0.003 |
| Drinking | 514(38.9) | 318(33.3) | 310(36.0) | 0.023 | 129(31.3) | 866(37.6) | 147(35.0) | 0.041 | 357(34.6) | 471(35.4) | 314(40.6) | 0.019 |
| PAL |  |  |  |  |  |  |  |  |  |  |  |  |
| Light PA level | 172(13.0) | 284(29.7) | 301(35.0) | <0.001 | 40(9.9) | 508(22.5) | 209(50.5) | <0.001 | 94(9.2) | 313(24.1) | 350(46.2) | <0.001 |
| Moderate PA level | 408(30.9) | 443(46.4) | 420(48.9) |  | 101(24.9) | 983(43.6) | 187(45.2) |  | 307(30.1) | 622(48.0) | 342(45.1) |  |
| Heavy PA level | 719(54.4) | 209(21.9) | 120(13.9) |  | 265(65.3) | 765(33.9) | 18(4.3) |  | 620(60.7) | 362(27.9) | 66(8.7) |  |
| TDEI (kcal) | 2515.2±545.7 | 2319.1±579.9 | 2329.6±557.0 | <0.001 | 2580.2±515.1 | 2412.9±562.3 | 2190.5±575.4 | <0.001 | 2524.7±546.0 | 2386.9±552.8 | 2265.7±582.3 | <0.001 |

<sup>a</sup> Data were expressed as numbers or percentages and presented as mean with standard deviation of the mean. Non-normally distributed data like baseline age reported as median (IQR). All univariate comparisons across subgroups were evaluated using ANOVA, Chi-square test, and Kruskal-Wallis as appropriate.
BMI: body mass index; PAL: physical activity level; TDEI: total daily energy intake; the source data can be found in Supplementary files-source data 1.

The age of participants with a high SES were higher than those with a low SES (p < 0.001). Compared to participants with middle and high SES, the participants with low SES were likely to reside in rural areas. With regards to lifestyles behaviors, the total daily energy intake and the percentage of smoking progressively decreased with the increment of SES. In the low SES group, the participants had more heavy physical activities than those in other two SES groups (p < 0.001). The proportion of drinking was higher among participants with high education and occupational position compared to those with low education and occupational position.

According to three latent class BMI trajectories, it was found that the total daily energy intake and the proportions of smoking and drinking were higher among participants in progressive obesity group, in comparison with other groups (Table S4). From normal-stable BMI group to progressive obesity group, heavy and moderate physical activities gradually decreased. In progressive obesity group, there were higher adult occupational position and parental occupational position, but no significant difference in participant’s education across the three trajectories (Figure 3).

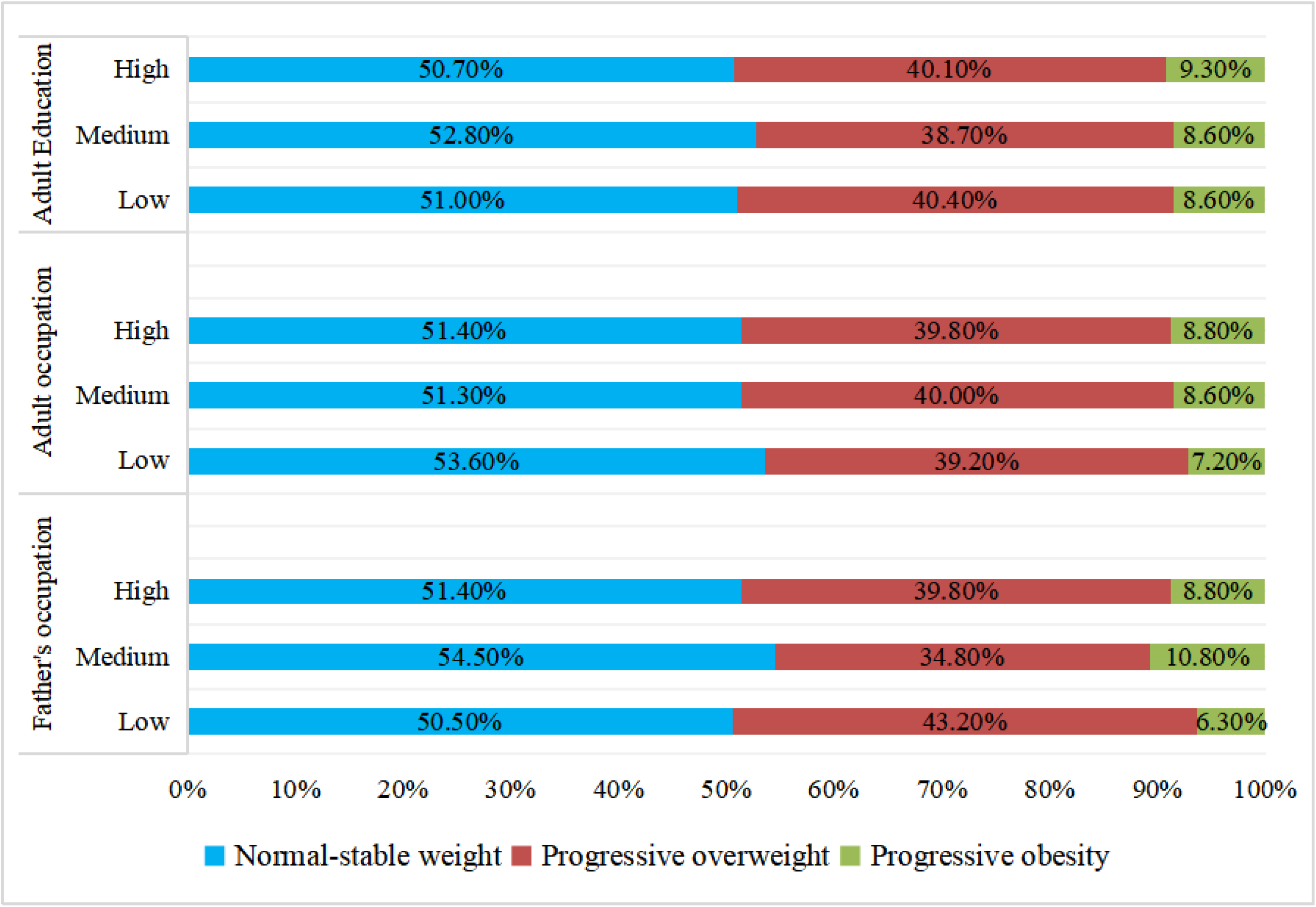

Through monomial logistic regression, participants with smoking (OR=1.35, 95% CI: 1.15-1.58; OR=1.53, 95% CI: 1.17-1.99) and drinking (OR=1.59, 95% CI: 1.22-2.06; OR=1.34, 95% CI: 1.16-1.57) were more likely to suffer from progressive obesity and overweight rather than normal-stable weight (Table 2). The females vs the males had much less risks for progressive obesity (OR=0.59, 95% CI: 0.44-0.77) and overweight (OR=0.62, 95%CI: 0.52-0.73). Participants in rural areas were more likely to develop into progressive overweight (OR=1.19, 95%CI: 1.02-1.40) in contrast with those in urban areas.

**Table 2.** The association of baseline characteristics with BMI trajectories^a^.

| Characteristics | Progressive obesity vs<br>normal-stable BMI | Progressive overweight vs<br>normal-stable BMI |
| --- | --- | --- |
|  | OR (95% CI) <sup>a</sup> | OR (95% CI) <sup>a</sup> |
| Gender |  |  |
| Male | 1.00(ref.) | 1.00(ref.) |
| Female | 0.59(0.44;0.77) | 0.62(0.52;0.73) |
| Place of residence |  |  |
| Urban | 1.00(ref.) | 1.00(ref.) |
| Rural | 0.95(0.73;1.24) | 1.19(1.02;1.40) |
| Smoking |  |  |
| Yes | 1.00(ref.) | 1.00(ref.) |
| No | 1.35(1.15;1.58) | 1.53(1.17;1.99) |
| Drinking |  |  |
| Yes | 1.00(ref.) | 1.00(ref.) |
| No | 1.59(1.22;2.06) | 1.34(1.16;1.57) |
| PAL |  |  |
| Light PA level | 1.00(ref.) | 1.00(ref.) |
| Moderate PA level | 0.80(0.59;1.10) | 1.22(1.01;1.48) |
| Heavy PA level | 0.74(0.53;1.04) | 1.16(0.95;1.42) |
| TDEI | 1.00(1.00;1.00) | 1.00(1.00;1.00) |

<sup>a</sup>Data were expressed as OR and 95% CI, using multinomial logistic regression model adjusting for all socioeconomic status in early life and adult life.
OR: odds ratios; CI: confidence interval; PAL: physical activity level; TDEI: total daily energy intake; the source data can be found in Supplementary files-source data 1.

### Association of socioeconomic status in early life and adult life with BMI trajectories

Table 3 showed that participants with the highest education attainment (OR=1.57, 95% CI:1.05, 2.35) or the highest adult occupational position (OR=1.17, 95% CI:1.01, 1.36), were more likely to have progressive obesity development than those with the lowest education attainment or adult occupational position. High father’s occupation vs low father’s occupation (OR=1.91, 95% CI:1.36, 2.69) was more likely to contribute to progressive obesity compared to normal-stable BMI. Similarly, apart from father’s occupation, high participant education (OR=1.33, 95% CI:1.05, 1.69) and adult occupational position (OR=1.20, 95% CI:1.10, 1.31) were more likely to increase the risk for progressive obesity in comparison with low participant education and adult occupational position after adjustment for all potential confounders. In the study, multiple adjustments resulted in some changes of these estimates but the pattern of association persisted (Table 3). We also found that the risk of obesity was lower in the smoking group and drinking increased the risk of obesity, compared to the normal-stable BMI (Figure S2 - S4, Supplementary file 1)

**Table 3.** The association of socioeconomic status in early life and adult life with BMI trajectories.

|  | Progressive overweight vs normal-stable BMI |  |  |
| --- | --- | --- | --- |
|  | Highest vs Lowest father's occupational position | Highest vs Lowest adult occupation | Highest vs Lowest adult education |
|  | OR (95% CI) | OR (95% CI) | OR (95% CI) |
| Model 1: Gender+residence +age | 1.06(0.88;1.28) | 1.22(1.12;1.32) | 1.17(1.10;1.24) |
| Model 2: Model 1+smoking+drinking | 1.04(0.86;1.27) | 1.21(1.11;1.31) | 1.16(1.10;1.23) |
| Model 3: Model 1+PAL | 1.08(0.88;1.31) | 1.31(1.03;1.64) | 1.22(1.14;1.30) |
| Model 4: Model 1+TDEI | 1.07(0.88;1.29) | 1.23(1.16;1.31) | 1.18(1.12;1.26) |
| model 5: Model 1+all risks | 1.06(0.86;1.30) | 1.33(1.05;1.69) | 1.20(1.10;1.31) |
|  | Progressive obesity vs normal-stable BMI |  |  |
|  | Highest vs Lowest father's occupational position | Highest vs Lowest adult occupation | Highest vs Lowest adult education |
|  | OR (95% CI) | OR (95% CI) | OR (95% CI) |
| Model 1: Gender+residence +age | 1.90(1.36;2.67) | 1.78(1.25;2.54) | 1.27(1.10;1.45) |
| Model 2: Model 1+smoking+drinking | 1.88(1.34;2.63) | 1.75(1.23;2.50) | 1.25(1.09;1.43) |
| Model 3: Model 1+PAL | 1.84(1.28;2.63) | 1.57(1.06;2.32) | 1.14(1.02;1.27) |
| Model 4: Model 1+TDEI | 1.93(1.76;2.13) | 1.79(1.25;2.56) | 1.27(1.11;1.45) |
| model 5: Model 1+all risks | 1.91(1.36;2.69) | 1.57(1.05;2.35) | 1.17(1.01;1.36) |

Data were expressed as OR and 95% CI, using multinomial logistic regression model adjusting for some confounders.
OR: odds ratios; CI: confidence interval; PAL: physical activity level; TDEI: total daily energy intake; the source data can be found in Supplementary files-source data 1.

### The association of life-course socioeconomic trajectories with BMI trajectories

Two socioeconomic indicators were combined to represent the socioeconomic trajectory from childhood to adulthood, including stable high, downward, upward, stable low groups. Table 4 presented the association of life-course socioeconomic trajectories with BMI trajectories. For progressive overweight vs normal-stable BMI, the odds ratio of participants with low SES in childhood but high SES in adulthood relative to a stable-low SES trajectory (low SES in both childhood and adulthood) was 1.56 (95% CI: 1.45, 1.68) after adjustment for potential confounders. Participants who were socially downwardly mobile (high SES in childhood and low SES in adulthood) or stable high in both childhood and adulthood, had a 1.26 and a 1.40-fold increased risk of progressive overweight, respectively, compared to those who had a stable-low SES trajectory. Physical activity explained 6.17% to 14.53% of the association between life-course socioeconomic trajectories with progressive obesity trajectory, TDEI explained 1.50 % to 1.95% of the association, and all risk factors explained 2.09% to 7.55% of the association.

**Table 4.** The association of life-course socioeconomic trajectories with BMI trajectories ^a^.

| Progressive overweight vs<br>normal-stable BMI | Stable low | upward |  | downward |  | Stable high |  |
| --- | --- | --- | --- | --- | --- | --- | --- |
| | OR (95% CI) | OR (95% CI) | %$\Delta^b$ (95% CI) | OR (95% CI) | %$\Delta$ (95% CI) | OR (95% CI) | %$\Delta$ (95% CI) |
| Model 1: Gender+residence+age | 1.00(ref.) | 1.51(1.41;1.62) | (ref.) | 1.22(1.10;1.35) | (ref.) | 1.32(1.24;1.41) | (ref.) |
| Model 2: Model 1+smoking+drinking | 1.00(ref.) | 1.51(1.41;1.62) | 0.01(-0.31;0.35) | 1.22(1.10;1.35) | 0.12(-2.90;3.15) | 1.32(1.24;1.41) | -0.48(-1.83;0.88) |
| Model 3: Model 1+PAL | 1.00(ref.) | 1.54(1.43;1.65) | 4.49(1.27;7.71) | 1.25(1.13;1.38) | 10.55(5.07;16.03) | 1.35(1.26;1.45) | 7.94(3.93;11.95) |
| Model 4: Model 1+TDEI | 1.00(ref.) | 1.52(1.41;1.63) | 0.86(-0.35;2.10) | 1.24(1.12;1.37) | 6.93(3.68;10.19) | 1.34(1.26;1.43) | 4.14(1.81;6.48) |
| Model 5: Model 1+all risks | 1.00(ref.) | 1.56(1.45;1.68) | 8.19(4.59;11.79) | 1.26(1.13;1.39) | 13.66(3.98;23.3) | 1.40(1.30;1.50) | 19.42(6.56;32.29) |
| Progressive obesity vs<br>normal-stable BMI | Stable low | upward |  | downward |  | Stable high |  |
| | OR (95% CI) | OR (95% CI) | %$\Delta$ (95% CI) | OR (95% CI) | %$\Delta$ (95% CI) | OR (95% CI) | %$\Delta$ (95% CI) |
| Model 1: Gender+residence+age | 1.00(ref.) | 1.29(1.11;1.48) | (ref.) | 1.95(1.63;2.33) | (ref.) | 2.45(2.18;2.76) | (ref.) |
| Model 2: Model 1+smoking+drinking | 1.00(ref.) | 1.29(1.11;1.49) | 0.72(-1.00;2.43) | 1.98(1.66;2.37) | 2.45(0.55;4.35) | 2.46(2.18;2.77) | 0.43(-0.33;1.20) |
| Model 3: Model 1+PAL | 1.00(ref.) | 1.15(1.00;1.34) | -14.53(-23.49;-5.47) | 1.87(1.57;2.24) | -6.17(-9.86;-2.50) | 2.17(1.91;2.47) | -13.47(-19.66;-7.29) |
| Model 4: Model 1+TDEI | 1.00(ref.) | 1.31(1.13;1.52) | 2.47(1.59;3.35) | 1.95(1.63;2.33) | -0.24(-2.34;1.86) | 2.48(2.21;2.80) | 1.50(0.30;2.68) |
| Model 5: Model 1+all risks | 1.00(ref.) | 1.21(1.04;1.41) | -7.55(-13.85;-1.25) | 1.93(1.61;2.31) | -2.09(-3.91;-0.27) | 2.35(2.06;2.67) | -4.75(-9.05;-0.45) |

<sup>a</sup> Life-course socioeconomic trajectory refers to father's occupational position and participants' occupational position.
<sup>b</sup> $\Delta$ , attenuation, representing the proportion of the SES–BMI trajectory association explained by the risk factor in question. % attenuation is calculated only for statistically significant associations.
OR: odds ratios; CI: confidence interval; PAL: physical activity level; TDEI: total daily energy intake; the source data can be found in Supplementary files-source data 1.

The risks for progressive obesity trajectory in participants with stable-high SES trajectory (OR=2.35, 95% CI: 2.06-2.67) was higher than that in participants with stable-low SES trajectory. Participants in downwardly SES trajectory had 1.93 times (95% CI: 1.61-2.31) higher risk for progressive obesity, and participants in upwardly SES trajectory had a 1.21 (95% CI: 1.04-1.41) times risk of progressive obesity, compared to the stable-low SES trajectory. Physical activity explained 4.49% to 10.55% of this increased risk for progressive obesity with life-course SES trajectories, TDEI explained 4.14% to 6.93% of the association, and all risk factors explained 8.19% to 19.42% of the association.

### The association of cumulative socioeconomic score with BMI trajectories

Table 5 showed that individuals with the highest life-course cumulative SES compared to those with the lowest life-course cumulative SES score, were 2.01 times (95% CI: 1.12; 4.00) more likely to be in progressive obesity trajectory. Physical activity (9.9%) could explain the highest proportion of this gradient. All risk factors combined attenuated by 1.8% for the OR of progressive obesity in the highest vs the lowest cumulative score groups. However, no significant associations between life-course cumulative SES and progressive overweight were found.

**Table 5.**
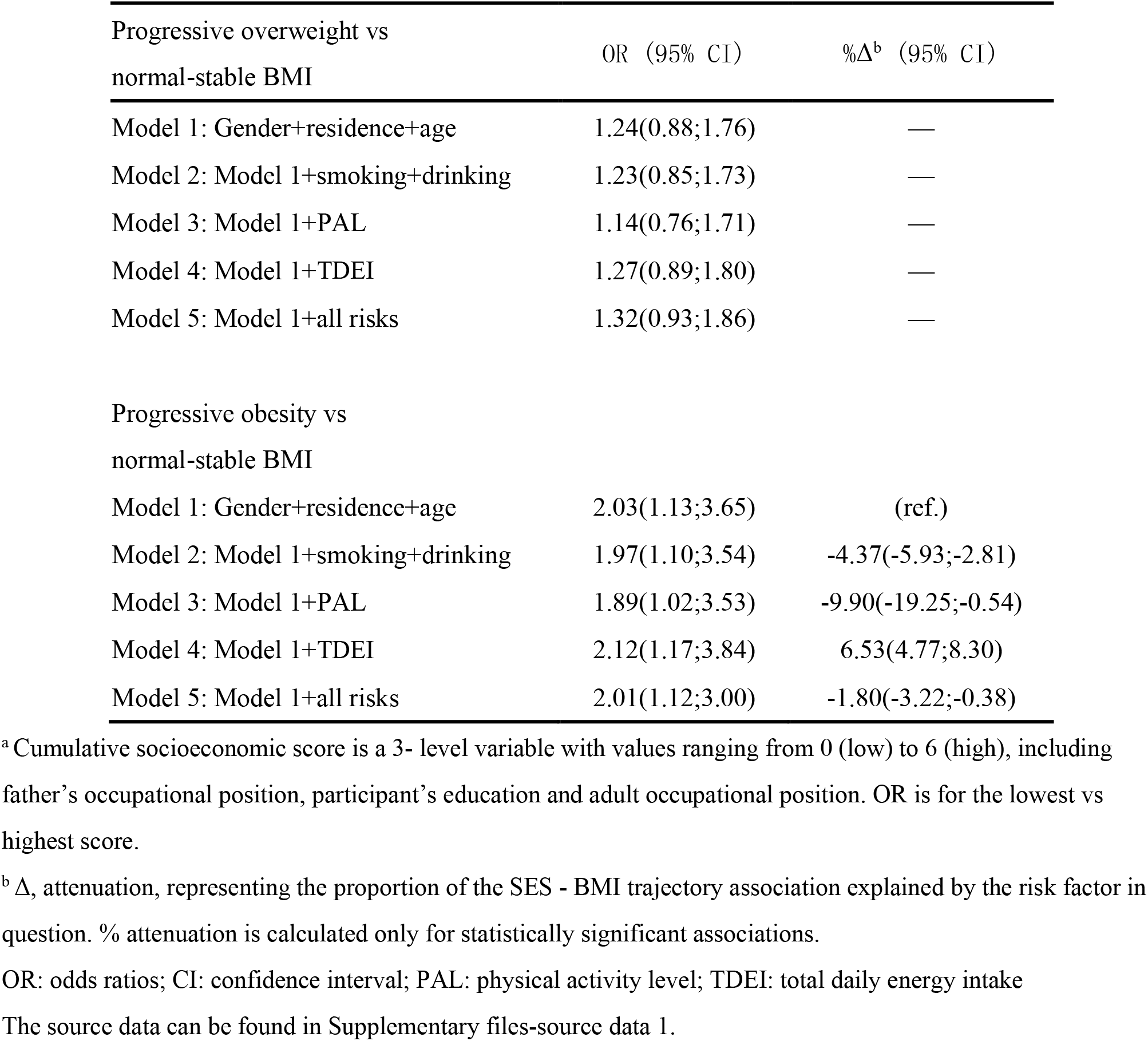
The association of cumulative socioeconomic score with BMI trajectories^a^.

### Sensitivity Analyses

After excluding 121 participants who developed diabetes, hypertension, myocardial infarction, stroke or cancer at baseline (Table S5), we had a total of 3017 participants. The results indicated that association of socioeconomic status in early life and adult life with BMI trajectories did not significantly change before and after excluding chronic diseases (Figure S2 - S4). There is no difference in the association of socioeconomic status in early life and adult life with BMI trajectories before and after multiple imputation, suggesting the robustness of the results (Figure S5 - S7).

## DISCUSSION

In the study, we built two indicators of life-course social class: a measure of social trajectories from childhood to adulthood and a cumulative score of individual socioeconomic indicators through life time. Our study established the BMI development trajectory in adulthood to reflect the dynamic change, and identified three patterns of the BMI trajectories among Chinese adults, including normal-stable BMI, progressive overweight, progressive obesity. High SES in early and adult life contributed to the increased long-term risk of obesity in Chinese adults. We explored the relationship between life-course SES and BMI development trajectory in China. Social disparities in BMI over the adult life course, may increase the future health risks associated with obesity in these populations as they age. Therefore, targeted interventions for these groups, especially early in the life course, have important public health implications.

Our study found that the hazards of the obesity increased with father’s occupational position in early life, participants’ occupational position and educational attainment in adult life. Higher SESs in developed countries generally had lower hazards of obesity(9, 22), but this pattern does not apply to China. Low SES limited the opportunities available for excess to food consumption and increases physical labor, while high SES increased access to excess food and avoids physical labor. These conditions contribute to the differences in weight gain between low SES groups and wealthy individuals in developing countries. Studies have showed that socioeconomic status in early life and perhaps even in earlier generations have significant influence on adult obesity development(23–25). However, this relationship still varies across countries with different levels of development. A cohort study from China reported a positive association between childhood SES and waist circumference in adulthood among the males(26). Low SES in childhood tended to have reduced risk of general obesity in adulthood among Hong Kong Chinese adults(27). Our findings were consistent with previous studies. There are several explanations for how exposure to low SES in childhood influence health later in life. Firstly, in different developmental stages, the impact of SES-related factors on health varied, with the greatest effects in specific stages (e.g. in utero, early childhood, adolescence). A cohort study US showed that in the prenatal period and the first year of life, the SES had significant effect on adult BMI but not in other periods of childhood(28). Secondly, the effect of poor SES in early life on adult health varied with the intensity and duration of exposure to socioeconomic disadvantage. Studies found that a longer duration of exposure to early childhood poverty would experience accelerated BMI growth trajectories in future(29).

The life-course socioeconomic trajectory reflects the intergenerational mobility of socioeconomic status between parents and children, from a life-course perspective, drawing attention to the powerful connections between individual lives and the historical and socioeconomic contexts of those live(30). Our results showed the individuals with higher life course SES, had a higher risk for long term obesity trend compared with individuals with stable-low SES. Upwardly mobile participants were more likely to fall into obesity than participants remaining in stable low social class. The associations observed in our study were consistent with previous studies in low-middle income countries showing positive correlation between life-course SES and obesity(31, 32). For example, a cross-sectional study in five middle-income countries (China, South Africa, India, Russia, and Mexico) found that life-course stable high or declining SES was associated with increased risks of overweight/obesity(31). In addition, our results contradicted findings in some previous studies conducted mostly in developed countries(33–35). For example, Albrecht and Gordon-Larsen reported upward education mobility was associated with low adult mean BMI(33); Sinead and Timothy found downwardly SES mobility were significantly more likely to be overweight/obese compared with those who remained of high SES(34). Thus, the relationship between life course SES and obesity seems to vary depending on the culture and socioeconomic development stage of the sample. Furthermore, those whose parents have high SES, may have the resources to increase consumption of fast food, high-sodium diets, and high-calorie beverages, which are thought to be significant dietary changes that promote obesity(36).

The accumulation model proposes that adverse exposures over the life course cause cumulative damage to biological systems. A positive association between cumulative socioeconomic score and obesity was found in our study, although some studies of Chinese populations did not find significant differences in obesity between groups with the greatest and the least cumulative disadvantage(27, 37). According to previous researches, a positive association with SES and obesity was shown in adults(10, 31), but the future burden of overweight and obesity may shift to populations with lower socioeconomic status. Further research is still needed to clarify the relationship between the socioeconomic transition occurring in China and obesity and related chronic diseases in adulthood.

Moreover, the potential mechanisms explaining the association between life-course SES and overweight/obesity also include socially patterned behavioral factors, such as alcohol use, dietary quality and physical activity(20). Current alcohol drinking was associated with obesity in our analyses. Alcohol consumption has a greater acute effect on calorie intake than other lifestyle factors(38). Furthermore, metabolic evidence also suggests that alcohol consumption may be associated not only with a higher urge to drink, but also with less restrictive eating behaviors, which further cause higher BMI(39). Due to the economic transition and introduction of the western lifestyle in China, people’s work and dietary patterns have changed dramatically. According to our analysis, people of low socioeconomic status are more likely to be engaged in moderate and heavy work, although they have a higher average daily energy intake compared to people of high socioeconomic status.

Considering the study data from a 20-year follow-up cohort in China, we are able to clearly confirm the association between life-course socioeconomic status and the risk of overweight/obesity. Additionally, we constructed the BMI development trajectories, which was a more stable and reliable indicator to reflect the long term BMI change. In the meanwhile, the two measures of life-course SES (life-course socioeconomic trajectories and a cumulative SES index) were also adopted to explore their origins of SES disparities in early life and long term effect of SES on obesity.

In this study, some limitations should be noted in the explanation of the results. Firstly, the socioeconomic level and lifestyle were mainly self-reported, and only baseline information was used, thus recall bias was inevitable. Second, because of lacking of parents’ income, education and other related SES indicators to represent the SES in early life of participants, the life course SES variable was constructed only using on paternal occupation and personal occupation position. Future studies with more reliable indicators in early life are preferred. Third, we excluded those with major chronic diseases at baseline, and then got robust results. However, the possibility of reverse causation and residual confounding may still exist in our study, due to many other unmeasured diseases. Fourth, our results may be subjected to other potential confounders that were not observed in the study.

### Conclusion

This study confirmed that socioeconomic status played an important role in the development of overweight and obesity. These patterns suggest that the effect of SES on adult BMI may act during critical periods in early life and accumulate throughout the lifespan. Physical activity and total daily energy intake partially explain this gradient, showing that both play an important role in obesity interventions. When implementing prevention measures of obesity in targeted groups, attention should be paid to the high socioeconomic status, especially in early life, such as childhood, adolescence.

## Access to research materials/Data sharing

China Health and Nutrition Survey data are available in a public, open access repository. Original data are available in Carolina Population Center of the University of North Carolina at Chapel Hill (https://www.cpc.unc.edu/projects/china).

## Additional files

Supplementary files

- Supplementary file 1. Table S1 to Table S6 and Figure S1 to Figure S7
- Source data 1. Sample analysis data for Table 1 to Table 4 and Figure 3
- Source data 2. Adult BMI trajectories data for Figure 2
- Source code. Analysis code based on Stata and R software
- Transparent reporting form

## SUPPLEMENTARY MATERIALS

**Table S1.** Social class based on self-reported primary occupation.

| **Social class** | **Primary occupation** |
| --- | --- |
| Class I | 1. Senior professional/technical worker |
| Class II | 1. Junior professional/technical worker<br>2. Administrator/executive/manager<br>3. Army officer, police officer |
| Class III-M (skilled manual) | 1. Skilled worker |
| Class III-NM (skilled non-manual) | 2. Office staff<br>3. Soldier, policeman<br>4. Athlete, actor, musician |
| Class IV (semi-skilled) | 1. Driver<br>2. Service worker |
| Class V (unskilled) | 1. Farmer, fisherman, hunter<br>2. Non-skilled worker |

**Table S2.** Definition of life course socioeconomic trajectories.

| Father's occupational position | Adult occupational position |  |
| --- | --- | --- |
|  | Low | High |
| Low | stable low | upward |
| High | downward | stable high |

**Table S3.** Tabulated Bayesian Information Criterion (BIC) for all participants.

| Number of groups | BIC<sup>1</sup> | BIC<sup>2</sup> | AIC | G<sup>2</sup> | Entropy | Group Membership Probabilities |
| --- | --- | --- | --- | --- | --- | --- |
| 1 | -29209.11 | -29207.17 | -29198.09 | -29195.09 | - | (100.00%) |
| 2 | -27113.79 | -27109.26 | -27088.08 | -27081.08 | 0.759 | (70.91%/29.08%) |
| 3 | -26325.60 | -26318.49 | -26285.21 | -26274.21 | 0.737 | (51.40%/39.80%/8.80%) |
| 4 | -26023.02 | -26012.67 | -25964.26 | -25948.26 | 0.677 | (28.79%/41.79%/23.37%/6.03%) |
| 5 | -25879.80 | -25866.86 | -25806.35 | -25786.35 | 0.714 | (41.50%/26.43%/24.42%/6.62%/1.01%) |

BIC<sup>1</sup>=Bayesian information criterion (for the total number of observations).
BIC<sup>2</sup>=Bayesian information criterion (for the total number of participants).
Abbreviations: BIC, Bayesian information criterion; AIC, Akaike information criterion.

**Table S4.** Participant characteristics across BMI trajectory groups (n = 3138)

|  | Normal-stable BMI | Progressive overweight | Progressive obesity |  |
| --- | --- | --- | --- | --- |
| Characteristics |  |  |  | *p* |
| participants(n) | 1613(51.4) | 1249(39.8) | 276(8.8) |  |
| Baseline age(years) | 22.0(6.0) | 23.0(9.0) | 25.00(9.0) | <0.001 |
| Gender |  |  |  |  |
| Male | 1068(66.2) | 949(76.0) | 212(76.8) | <0.001 |
| Female | 545(33.8) | 300(24.0) | 64(23.2) |  |
| Residential areas |  |  |  |  |
| Urban | 573(35.5) | 394(31.5) | 101(36.6) | 0.054 |
| Rural | 1040(64.5) | 855(68.5) | 175(63.4) |  |
| BMI(Kg/m<sup>2</sup>) | 19.6±1.5 | 22.1±2.1 | 25.8±3.2 | <0.001 |
| *Socioeconomic indicators* |  |  |  |  |
| Father's occupational position |  |  |  |  |
| Low | 668(41.4) | 571(45.7) | 83(30.1) | <0.001 |
| Medium | 520(32.2) | 332(26.6) | 103(37.3) |  |
| High | 425(26.3) | 346(27.7) | 90(32.6) |  |
| Adult occupation |  |  |  |  |
| Low | 554(34.3) | 405(32.4) | 74(26.8) | 0.029 |
| Medium | 683(42.3) | 533(42.7) | 115(41.7) |  |
| High | 376(23.3) | 311(24.9) | 87(31.5) |  |
| Adult education |  |  |  |  |
| Low | 373(23.1) | 298(23.9) | 63(22.8) | 0.843 |
| Medium | 733(45.4) | 546(43.7) | 119(43.1) |  |
| High | 507(31.4) | 405(32.4) | 94(34.1) |  |
| *Health-related behaviors* |  |  |  |  |
| Smoking | 467(29.0) | 442(35.4) | 106(38.4) | <0.001 |
| Drinking | 528(32.7) | 494(39.6) | 120(43.6) |  |
| PAL |  |  |  |  |
| Light PA level | 403(25.5) | 272(22.2) | 82(30.5) | 0.043 |
| Moderate PA level | 638(40.4) | 528(43.1) | 105(39.0) |  |
| Heavy PA level | 540(34.2) | 426(34.7) | 82(30.5) |  |
| TDEI (kcal) | 2348.6±660.4 | 2460.5±558.5 | 2479.±557.5 | <0.001 |

<sup>a</sup> Data were expressed as numbers or percentages and presented as mean with standard deviation of the mean. Non-normally distributed data like baseline age reported as median (IQR)
All univariate comparisons across subgroups were evaluated using ANOVA, Chi-square test, and Kruskal-Wallis as appropriate.
BMI: body mass index; PAL: physical activity level; TDEI: total daily energy intake

**Table S5.** Details about participants with chronic diseases at baseline(n=121)

| Chronic Disease | Number (%) |
| --- | --- |
| HYPERTENSION | 79(2.5) |
| DIABATES | 29(0.9) |
| STROKE | 7(0.2) |
| MYOCARDIAL INFARCTION | 5(0.2) |
| ASTHMA | 9(0.3) |
| CANCER | 6(0.2) |

Note: There were 14 participants with two or more chronic diseases in the total sample(n=3138).

**Table S6.** Details about missing covariaties.

| Variable | Missing ratio (%) |
| --- | --- |
| Smoking | 1 (0. 1) |
| Drinking | 3 (0. 1) |
| Physical activity level | 62 (2. 0) |
| Total daily energy intake | 77 (2. 5) |

**Figure S1.**
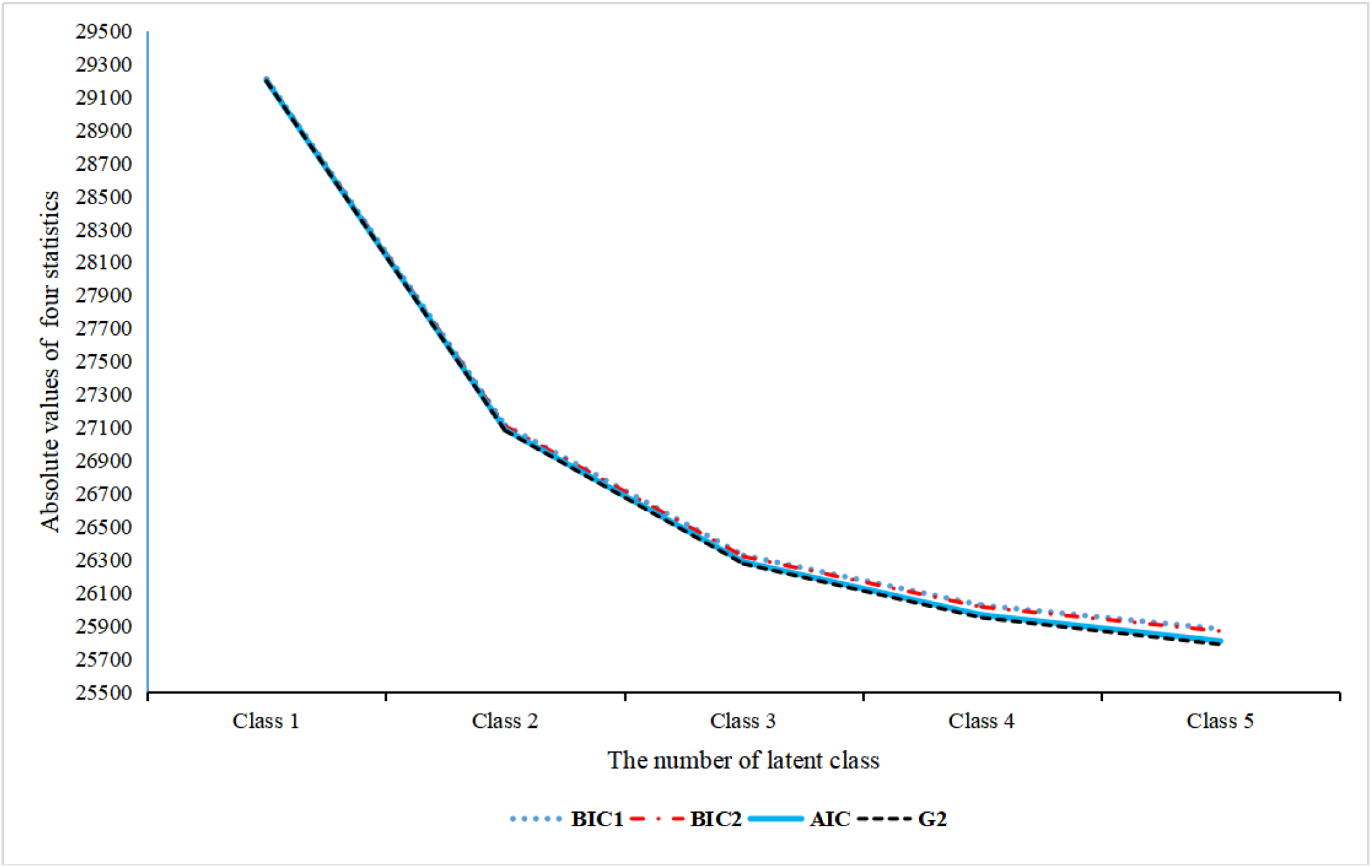
Model Fitting Indicators of Group-based trajectory model (GBTM)

**Figure S2.**
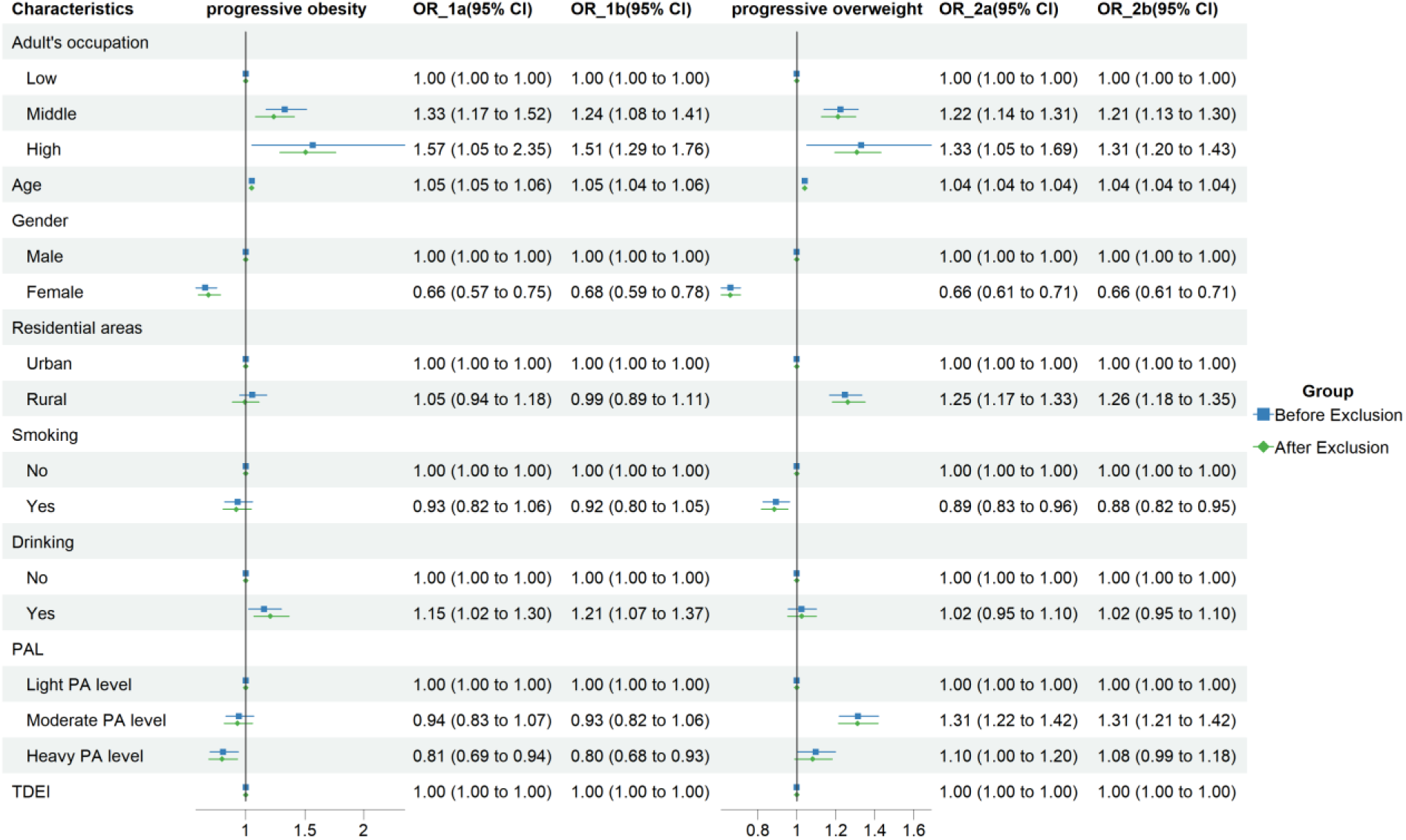
sensitivity analysis of the associations between BMI trajectories and adult’s occupation (before & after exclude chronic diseases) OR_1a: “progressive obesity” trajectory before excluding **chronic diseases;** OR_1b: “progressive obesity” trajectory after excluding **chronic diseases;** OR_2a: “progressive overweight” trajectory before excluding **chronic diseases;** OR_2b: “progressive overweight” trajectory after excluding **chronic diseases;** PAL: Physical Activity Level; TDEI: total daily energy intake

**Figure S3.**
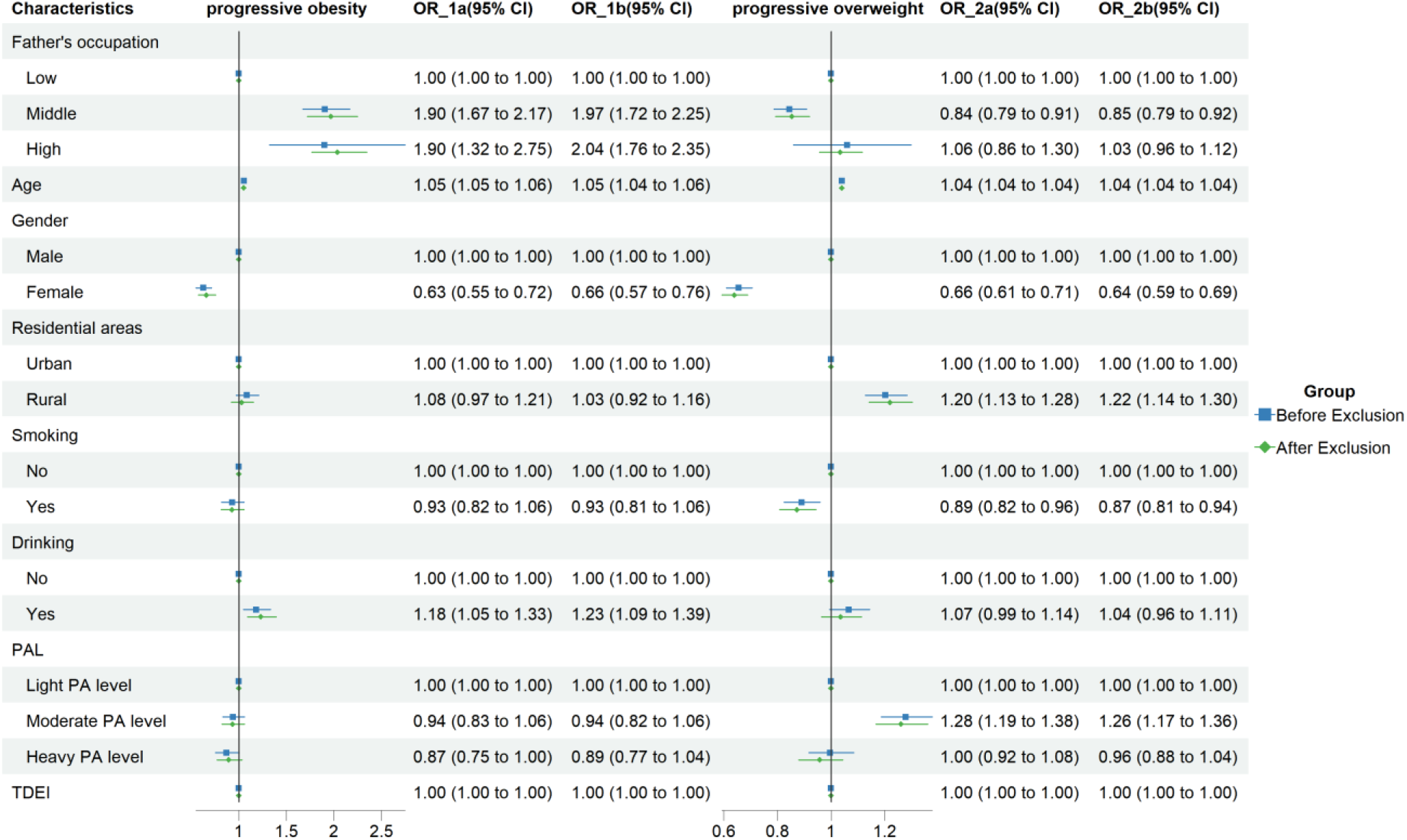
sensitivity analysis of the associations between BMI trajectories and father’s occupation (before & after exclude chronic diseases) OR_1a: “progressive obesity” trajectory before excluding **chronic diseases;** OR_1b: “progressive obesity” trajectory after excluding **chronic diseases;** OR_2a: “progressive overweight” trajectory before excluding **chronic diseases;** OR_2b: “progressive overweight” trajectory after excluding **chronic diseases;** PAL: Physical Activity Level; TDEI: total daily energy intake

**Figure S4.**
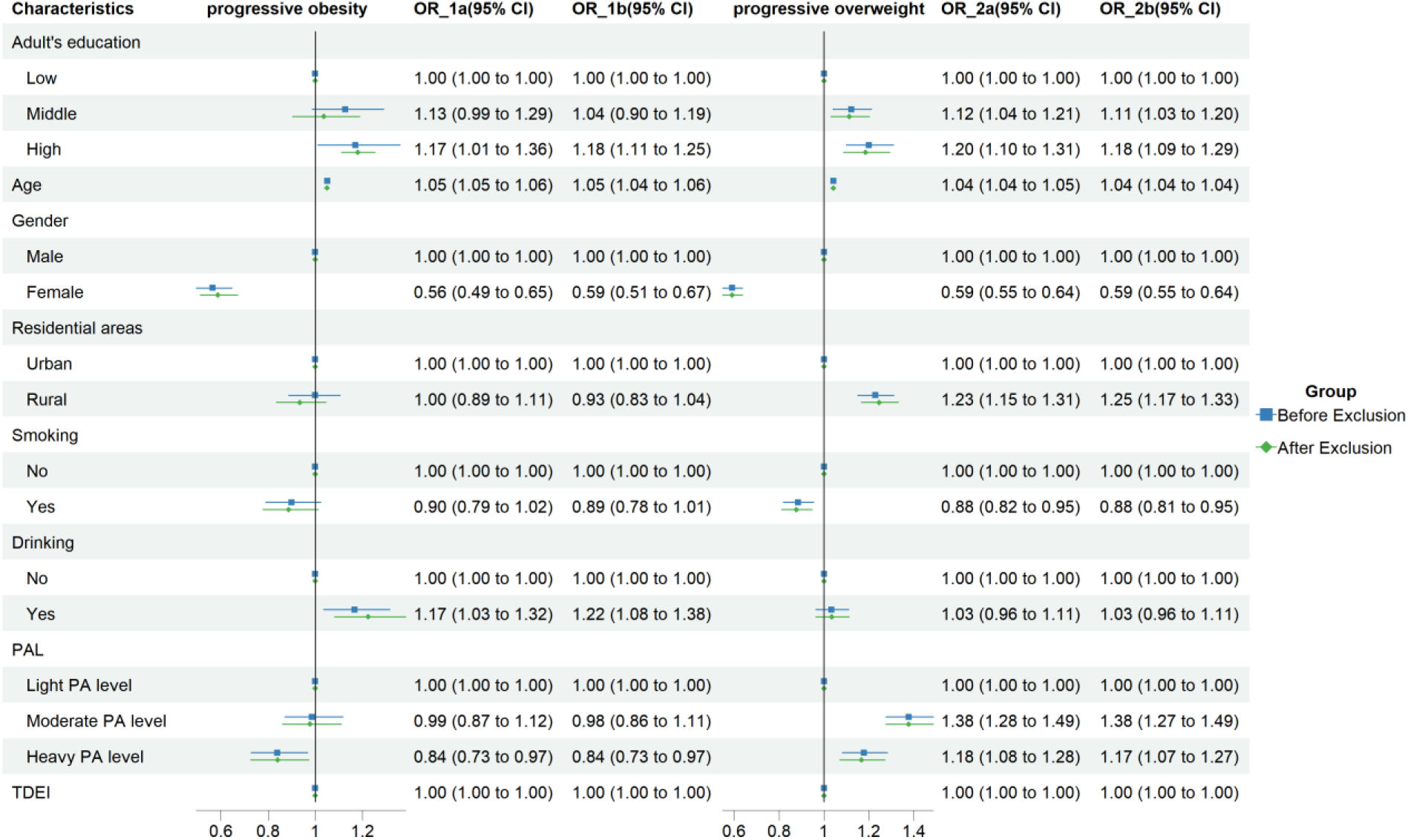
sensitivity analysis of the associations between BMI trajectories and adult’s education (before & after exclude chronic diseases) OR_1a: “progressive obesity” trajectory before excluding **chronic diseases;** OR_1b: “progressive obesity” trajectory after excluding **chronic diseases;** OR_2a: “progressive overweight” trajectory before excluding **chronic diseases;** OR_2b: “progressive overweight” trajectory after excluding **chronic diseases;** PAL: Physical Activity Level; TDEI: total daily energy intake

**Figure S5.**
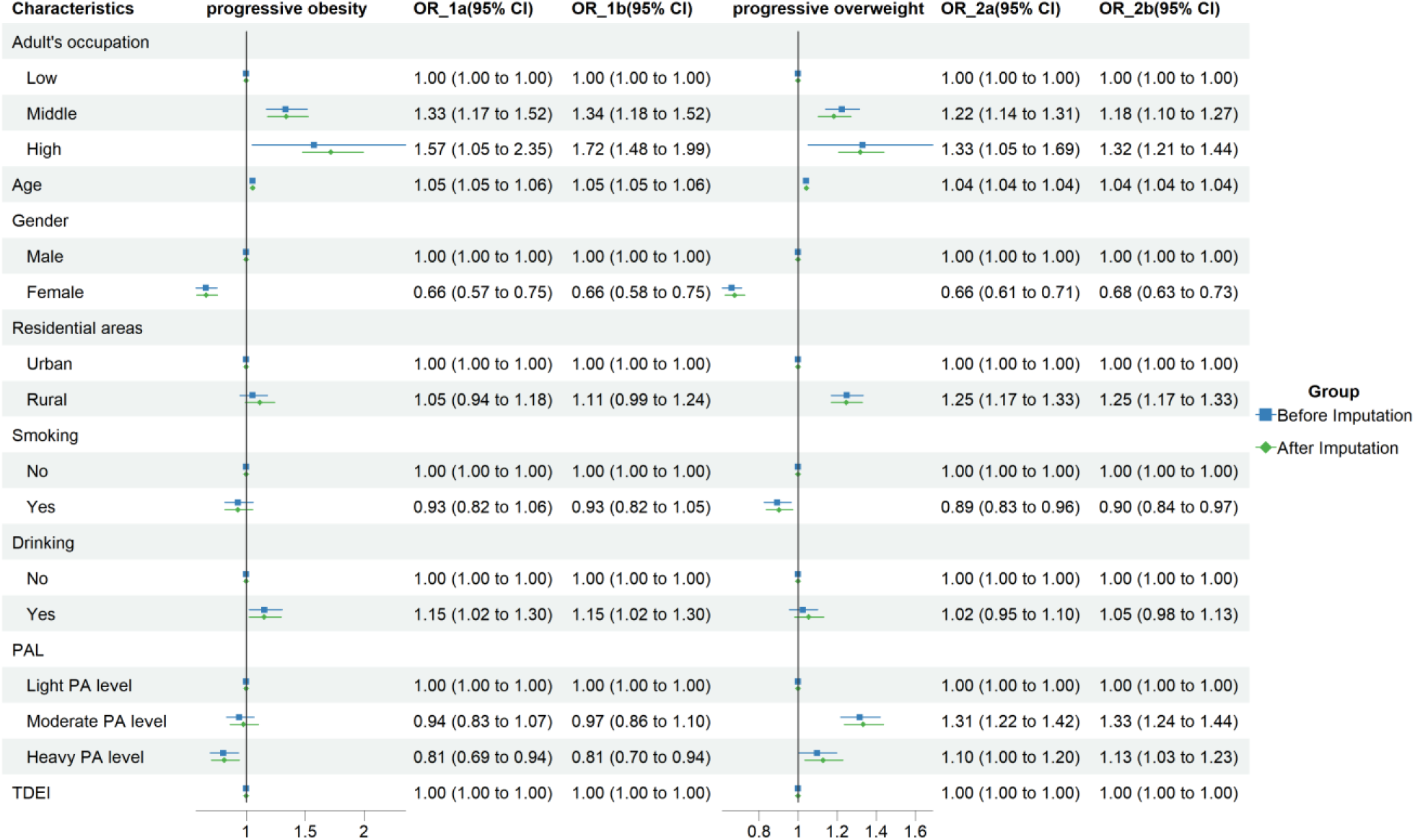
sensitivity analysis of the associations between BMI trajectories and adult’s occupation (before & after Multiple Imputation) OR_1a: “progressive obesity” trajectory before multiple imputation; OR_1b: “progressive obesity” trajectory after multiple imputation; OR_2a: “progressive overweight” trajectory before multiple imputation; OR_2b: “progressive overweight” trajectory after multiple imputation; PAL: Physical Activity Level; TDEI: total daily energy intake

**Figure S6.**
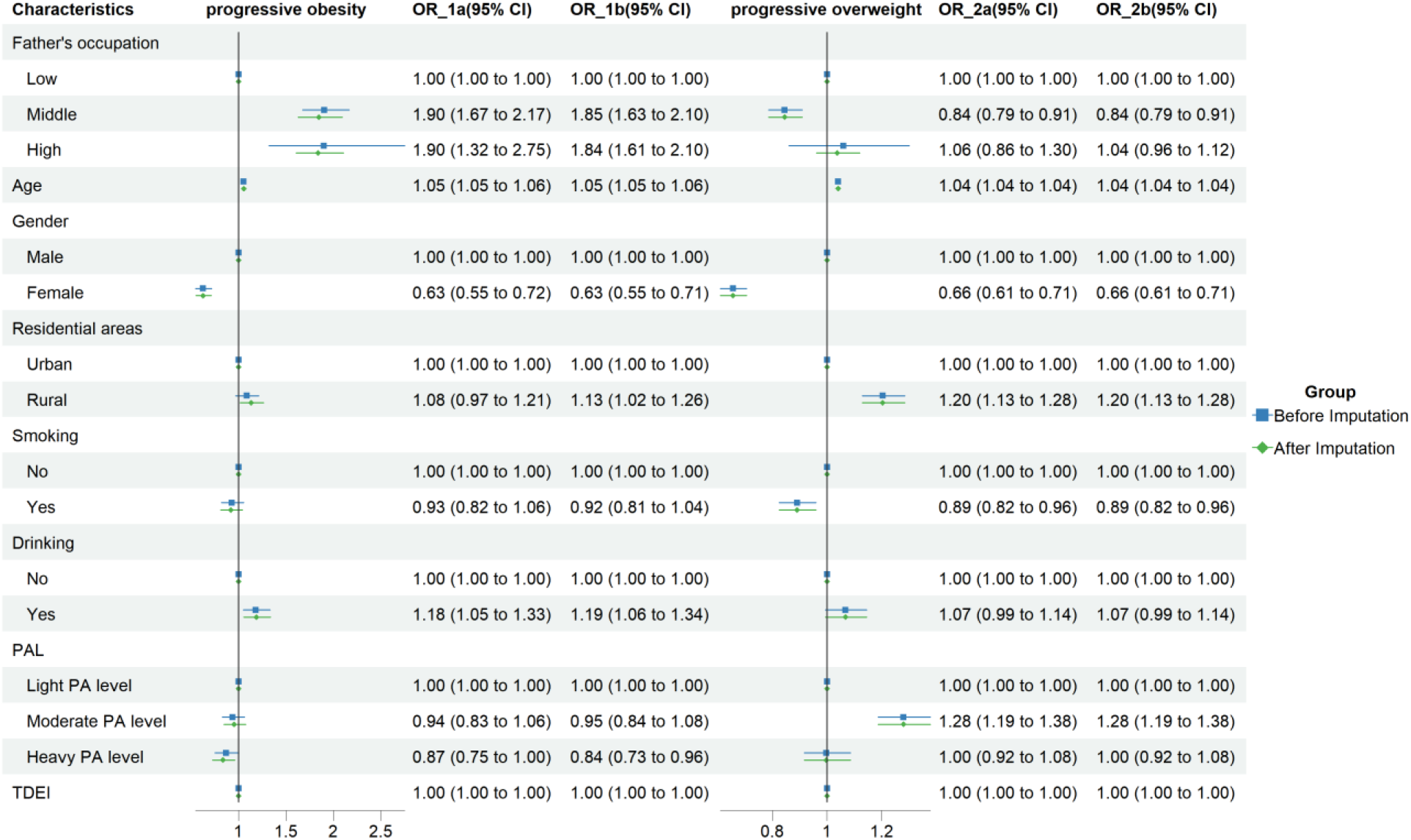
sensitivity analysis of the associations between BMI trajectories and father’s occupation (before & after Multiple Imputation) OR_1a: “progressive obesity” trajectory before multiple imputation; OR_1b: “progressive obesity” trajectory after multiple imputation; OR_2a: “progressive overweight” trajectory before multiple imputation; OR_2b: “progressive overweight” trajectory after multiple imputation; PAL: Physical Activity Level; TDEI: total daily energy intake

**Figure S7.**
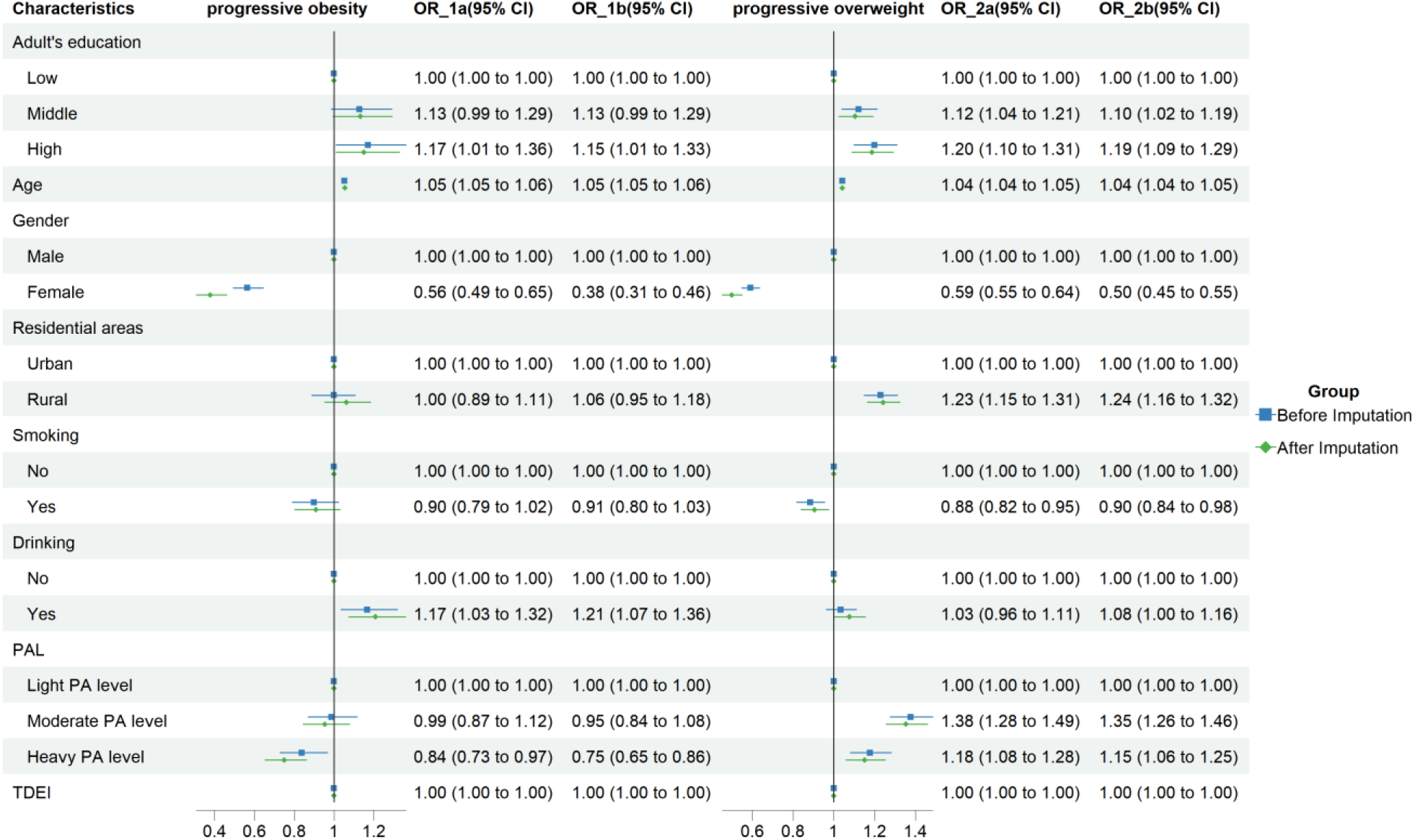
sensitivity analysis of the associations between BMI trajectories and adult’s education (before & after Multiple Imputation) OR_1a: “progressive obesity” trajectory before multiple imputation; OR_1b: “progressive obesity” trajectory after multiple imputation; OR_2a: “progressive overweight” trajectory before multiple imputation; OR_2b: “progressive overweight” trajectory after multiple imputation; PAL: Physical Activity Level; TDEI: total daily energy intake

